# Wildfires change plant scents but not pollinator attraction in a Mediterranean palm

**DOI:** 10.1101/2022.09.06.506788

**Authors:** Yedra García, María Clara Castellanos, Juli G. Pausas

## Abstract

Natural fire regimes are currently changing worldwide. These alterations may affect not only plant and animal species but also their interactions. Recently, a few studies have shown the effects of different disturbances on pollination through changes on plant fragrances mediating this interaction, yet no studies have focused on the effects of fires. Here, we assessed whether wildfires can modify plant scents and, in turn, pollinator attraction in a widespread palm in the western Mediterranean Basin. We studied the fire-adapted palm *Chamaerops humilis* and its nursery (dominant in unburnt sites) and non-nursery (dominant in recently burnt sites) beetle pollinators. In nursery pollination systems, where pollinators develop inside their host plant, volatile organic compounds (VOCs) emitted by plants to attract pollinators can be crucial because of the tight interdependence among the interacting species. However, these systems can also involve non-nursery copollinators whose importance is context dependent, and potentially relevant for plant success after disturbance. We first compare scent composition between plants growing in burned and unburned sites after recent wildfires; then we conducted olfactory bioassays with the two beetle pollinators. Fires changed the palm’s scent composition; however, the two pollinators responded similarly to scent from burnt and unburnt areas which may ensure plant reproduction even after recent fire events. We show, for the first time, that wildfires can alter plant fragrances mediating mutualistic interactions, and that flexible pollinator responses to variable odourscapes can enhance resilience in plant performance.

## Introduction

Wildfires are natural disturbances that play a significant role in many ecosystems, often shaping their ecology and evolution, from community composition to species adaptation (Pausas and Keeley, 2009). These natural fire regimes are currently changing, with a tendency towards increases in fire frequency and size in different regions (Westerling et al., 2006; Pausas and Fernández-Muñoz, 2012; Nicholson and Egan, 2019). The alteration of historical fire patterns may negatively impact species adapted to particular fire regimes (Keeley et al., 2011), and ultimately their interactions with other partners. For instance, changes in fire frequency can alter the ability of postfire recovery and resource allocation in plant species on which animals rely for food or reproduction after fires (Swab et al., 2012; Clarke et al., 2013). In the case of pollination, changes in fire frequencies can lead to a decline in pollinator populations or favour generalist species in detriment of pollinators with narrow habitat/food requirements (Lazarina et al., 2016; Carbone et al., 2019). Thus, assessing postfire pollination responses in fire-prone ecosystems is the first step in understanding how further changes on historical fire regimes will affect this key mutualism as it is plant pollination.

From the plant’s perspective, studies on disturbance effects on pollination have focused on plant visual communication (e.g. floral colour and shape) rather than chemical signals such as plant scents (Burkle and Runyon, 2017). Recently, a few studies have reported negative impacts of drought, ozone and CO_2_ emissions on pollination through the alteration of plant scents (e.g. Burkle and Runyon, 2016, 2017; Farré-Armengol et al., 2016; Glenny et al., 2018). Plant scents are chemical signals comprised of volatile organic compounds (VOCs) that mediate, among other things, plant-pollinator interactions, enhancing plant recognition and the attraction of pollinators even from long distances (Dudareva and Pichersky, 2006). These fragrances may attract a wide range of pollinator species (Dötterl et al., 2012) or only a few specialized pollinators (Proffit et al., 2009). Changes on plant scents (i.e., changes in scent amount and/or in volatile compound ratios that attract pollinators) can alter plant chemical signalling and ultimately disrupt plant-animal interactions (Farré-Armengol et al., 2016; Li et al., 2016). Wildfires may modify these scents by varying soil nutrient and water availability (Burkle and Runyon, 2017; Campbell et al., 2019; Luizzi et al., 2021). Moreover, fires may alter plant fragrances involved in pollination through postfire changes in herbivore pressures, as recent studies have shown that herbivores may also affect patterns of floral scent (Burkle and Runyon, 2017). However, to our knowledge, the impact of fire on pollination through changes in plant scents has not been addressed.

Here we explore the effects of wildfires on scents mediating plant-pollinator interactions in a widespread palm from the western Mediterranean Basin, *Chamaerops humilis* (Arecaceae). The palm is engaged in a nursery mutualism with the weevil *Derelomus chamaeropis* (Curculionidae), whose larvae grow inside the palm’s old inflorescences (Anstett, 1999; Dufaÿ and Anstett, 2004). Nursery pollination systems, where pollinators develop inside plants’ reproductive structures (Dufaÿ and Anstett, 2003), provide ideal examples to test for disturbance effects on scent-mediated interactions because of the tight relationship that is frequently mediated by chemical communication (Hossaert-McKey et al., 2010). During the flowering period, *C. humilis* emits a strong scent that attracts the pollinator to the new inflorescences (Dufaÿ et al., 2003). The palm can also have an effective non-nursery co-pollinator, the sap beetle *Meligethinus pallidulus* (Nitidulidae, García et al., 2018). Whether the scent also attracts the co-pollinator is unknown but expected as scent-mediated pollination is common in other Arecaceae with sap beetle pollinators (Knudsen et al., 2001).

After a wildfire, there is a marked reduction in the weevil’s abundance (in addition to a lack of food and sites for mating and breeding in the palm), and a temporary replacement by the quickly recolonizing sap beetle ensures the palm’s reproduction (García et al., 2018). That is, in the fire-prone landscapes where the plant is native, the dominance of each pollinator varies depending on the fire context: while the nursery (weevil) pollinator predominates in unburnt areas, the non-nursery co-pollinator (sap beetle) is dominant after recent fires (less than one-year after the fire). To assess whether wildfires change the plant’s fragrance, we collected and analysed the scent from adult plants in burnt and adjacent unburnt areas after recent wildfires in two localities in Eastern Spain. We then performed olfactory bioassays with the two beetle pollinators and scent collected in the field to test whether differences in pollinator attraction explained the variation in pollinator’s abundance after fires.

## METHODS

### Study system

*Chamaerops humilis* is a dwarf dioecious palm common in fire-prone shrublands of the western Mediterranean Basin. It sprouts quickly postfire from surviving apical buds, and can flower the spring following a fire that usually occurs in summer (Tavsanoglu and Pausas, 2018). Successful pollination depends on two pollen-feeding beetle species (Dufaÿ and Anstett, 2004; García et al., 2018). The larvae of the weevil *Derelomus chamaeropis* (Curculionidae) develop inside the palm’s old inflorescences during the winter, and adults emerge when the palm flowers (Anstett, 1999; Dufaÿ and Anstett, 2004). This interaction is mediated by the strong scent emitted by the leaves from both female and male palms during anthesis (Dufaÿ et al., 2003, 2004). Leaves emit much less scent before flowering, and the weak blend of the inflorescences does not attract the weevil (Dufaÿ et al., 2003, 2004; Caissard et al., 2004). Thus, despite being decoupled in space, the chemical signal (leaf scent) and the reward (pollen and brood sites in inflorescences) are temporally coupled during flowering.

The additional pollinator is the sap beetle *Meligethinus pallidulus* (Nitidulidae) which is smaller and carries less pollen, but it is an effective pollinator and relatively abundant in recent postfire conditions where the weevil is virtually absent (García et al., 2018). Contrary to the weevil, *M. pallidulus* does not develop inside the palm inflorescences but it is abundant on the stem and inflorescences during blooming.

### Scent collection and analysis

The study was carried out in 2017 during the flowering period (late March to May) at two sites in eastern Spain (Carcaixent and Xàbia, Fig. 1). The sites had been affected by high intensity wildfires in the previous summer (for more details on the study fires see García et al. 2018). At each site, *C. humilis* adult plants were studied inside the burnt and in an adjacent unburnt area (control; Fig. 1) coinciding with the first postfire flowering season. Sampled plants were separated from each other by at least 5 m, and from the fire’s perimeter by at least 50 m. We recorded the stem length of each sample plant with a measuring tape. To control for potential confounding effects of within-site variation in scent, we allowed sampled plants within the same study site to be closer to plants of the other treatment (e.g. a plant from the burnt and a plant from the unburnt) than to plants of the same treatment (e.g. two plants in the burnt). This means that a sampled plant from the burnt area could be closer to a sampled plant from the unburnt than to another plant from the burnt area within the same study site (e.g., Carcaixent). During scent collection, all sampled plants had inflorescences at anthesis, and collection took place at both sites during the morning coinciding with the period of activity of the two beetle species (Dufaÿ et al., 2003, 2004). Specifically, in Carcaixent we sampled 8 *C. humilis* individuals per sex at each burnt and control area (N=32 plants). The same was sampled in Xàbia but only 4 female plants were sampled within the burnt area (N=28 plants). We collected the scent in the field from a healthy leaf in each of the 60 plants using headspace adsorption. The leaf was enclosed in a polyethylene terephthalate bag (40 cm width x 49.5 cm length) for 10 minutes and then the volatiles were trapped for 5 minutes using scent traps connected by a Teflon tube to a 9V portable membrane pump (Appendix S1: Methods). We consider this collection time appropriate for *C. humilis* because of the strong scent emission that is even perceived by humans (Düfay et al. 2003). Other studies on floral scent have also collected plant fragrance during similar time periods (e.g. Prieto-Benítez et al. 2015, 2016). We collected ambient controls at each study site (one at each burnt and unburnt area) during 5 minutes with an empty bag opened at the bottom, to correct for contaminants (Appendix S1: Methods). Scent samples were analysed with gas chromatography-mass spectrometry in the Mass Spectrometry Section of The Experimental Research Support Service (SCSIE) of the University of Valencia (Appendix S1: Methods). Chromatogram peaks were integrated with MassHunter Software B.07.00 (Agilent). Identification of compound peaks was made by matching mass spectra to those of NIST 11 and Willey 9 libraries. To be conservative, only compounds with scores over 75 were included as identified while the rest of compounds remained as unknown. We built two scent data matrices with VOC information: i) presence or absence of VOCs, and ii) VOC proportions.

**Figure 1.**
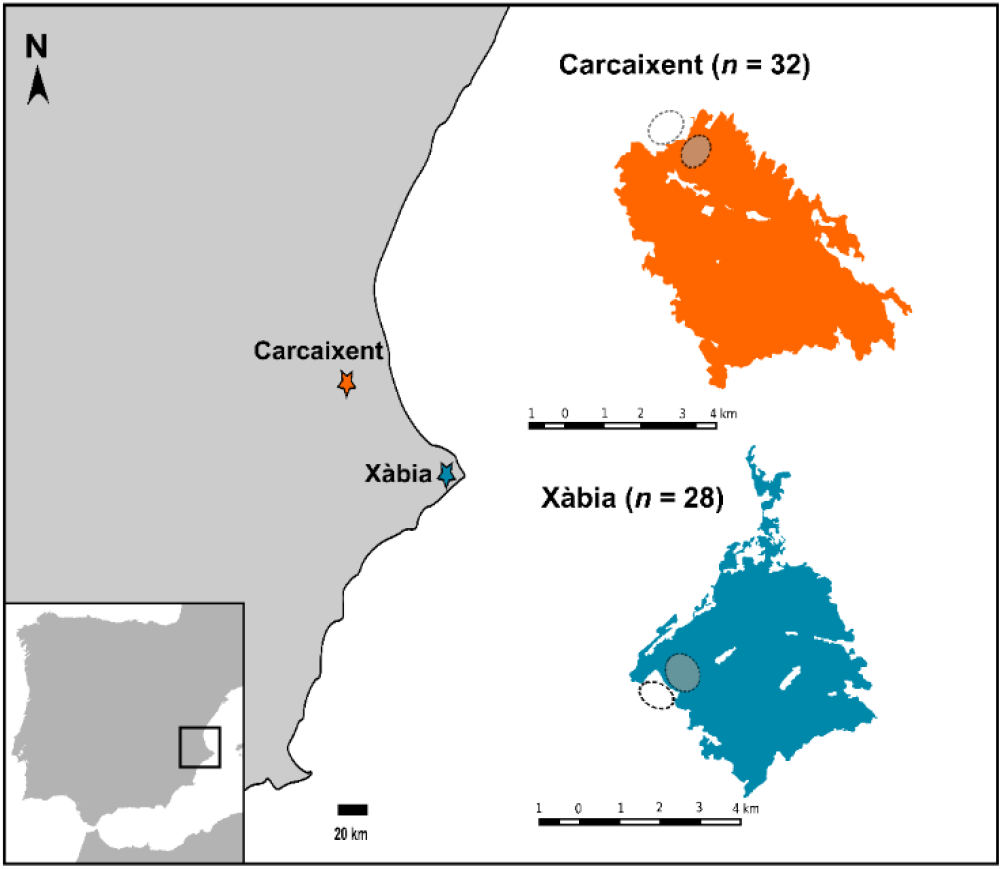
Location of two study sites, Carcaixent and Xàbia, in eastern Spain (left) and the fire perimeters (right) with the paired unburnt (dashed circles) and burnt (dashed grey circles) sampling areas.

### Olfactory bioassays

To test if the pollinators were attracted by *C. humilis* scent and whether the attractiveness of the signal differed between burnt and control (unburnt) areas, we ran olfactory bioassays using a glass Y-tube olfactometer as described in Dufaÿ et al. (2003, Appendix S1: Fig. S1). We sampled the leaf scent from four individuals per sex at burnt and unburnt areas that were different from those individuals used for floral scent comparisons. We followed a similar procedure as in Dufäy et al. (2003, Appendix S1: Methods), except that we used scent samples collected in the field to avoid potential scent changes after leaf cutting. Humidified clean air entered into the two arms through glass flasks containing a stripe of filter paper with i) 10 µl of *C. humilis* scent eluted in high-grade acetone (Chromasolv ®), or ii) the control with acetone only. For the bioassays, individuals of the two beetle species were collected from male inflorescences in unburnt and the surrounding areas of the two sites. We discarded to collect beetles in burnt sites, where beetle abundances where low, to not to alter the postfire recovery process. A single beetle individual was tested in the olfactometer at each trial. Each scent sample was tested over six trials alternating three individuals of each beetle species to ensure that the two pollinators were exposed to similar scents. The origin of the scent sample (study site) and plant sex were also alternated in the bioassays. To be able to compare beetles’ choice frequencies between treatments we alternated bioassays with control trials that had only acetone in the two arms of the olfactometer (as in Dufäy et al. 2003). We performed an extra series of bioassays to assess the specificity between the palm’s signal and the pollinators. To test the response specificity (i.e. the insect’s preference for the scent of a specific plant) of pollinators we collected flowers of 8 co-occurring species in the unburnt areas (*Cistus salvifolius, Genista scorpius, Gladiolus illyricus, Iris sisyrinchium, Lavandula angustifolia, Minuartia hybrida, Muscari neglectum* and *Rosmarinus officinalis*, Appendix S1: Methods). For this round of olfactory bioassays, the two beetle pollinators were collected from a third *C. humilis* population (Tivissa; 40^0^58’47” N, 0^0^41’35” E) that started flowering later. We gave 8 individuals of each beetle species the choice between scent from *C. humilis* and floral scent from one of eight co-occurring plant species (8 individual beetles tested for each pair of *C. humilis* and one co-occurring species, Appendix S1: Methods).

### Statistical analyses

To visualize the variation in scent composition between burnt and unburnt conditions, we defined the chemospace using a three-dimensional ordination space computed with nonmetric multidimensional scaling (NMDS) based on Bray-Curtis distances. The proportions of VOCs were fourth root transformed prior to ordination to reduce the influence of the most abundant compounds (Schlumpberger and Raguso, 2008). NMDS were run with the R package *vegan* (Dixon 2003).

To statistically assess whether fire induced changes on scent composition we used a model-based framework. Specifically, we fitted multivariate generalized linear models (MGLMs) with binomial (on the presence/absence scent data matrix and using the “cloglog” link to account for overdispersion) and tweedie (on the matrix with VOC proportions that shows a high number of zeros and using link “log” and variance power= 1.2) family distributions. The multivariate models included scent matrix: (i) presence/absence of each VOC or (ii) VOC proportions, as the response variable and fire treatment (unburnt vs. burnt), site, and their interaction as predictors (Appendix S1: Methods). To control for plant size effects that may influence scent amount, we included plant height (stem length in cm) as a covariate. Plant sex was not statistically significant in preliminary model runs (Fig. 2, Appendix S1: Table S2) and it was discarded from further MGLMs. Due to the statistical significance of study site, we fitted additional MGLMs to address the effect of fire on scent composition at each site. MGLMs were fitted with the packages *mvabund, statmot* and *tweedie* in R (Giner and Smyth, 2016; Dunn, 2017; Wang et al., 2017). To minimize within-site variation in scent composition (i.e. between burnt and unburnt areas of the same site) not related with the fires, we run the MGLMs on VOC proportions at each site with the subset of VOCs present in both areas (in burnt and unburnt at each site).

**Figure 2.**
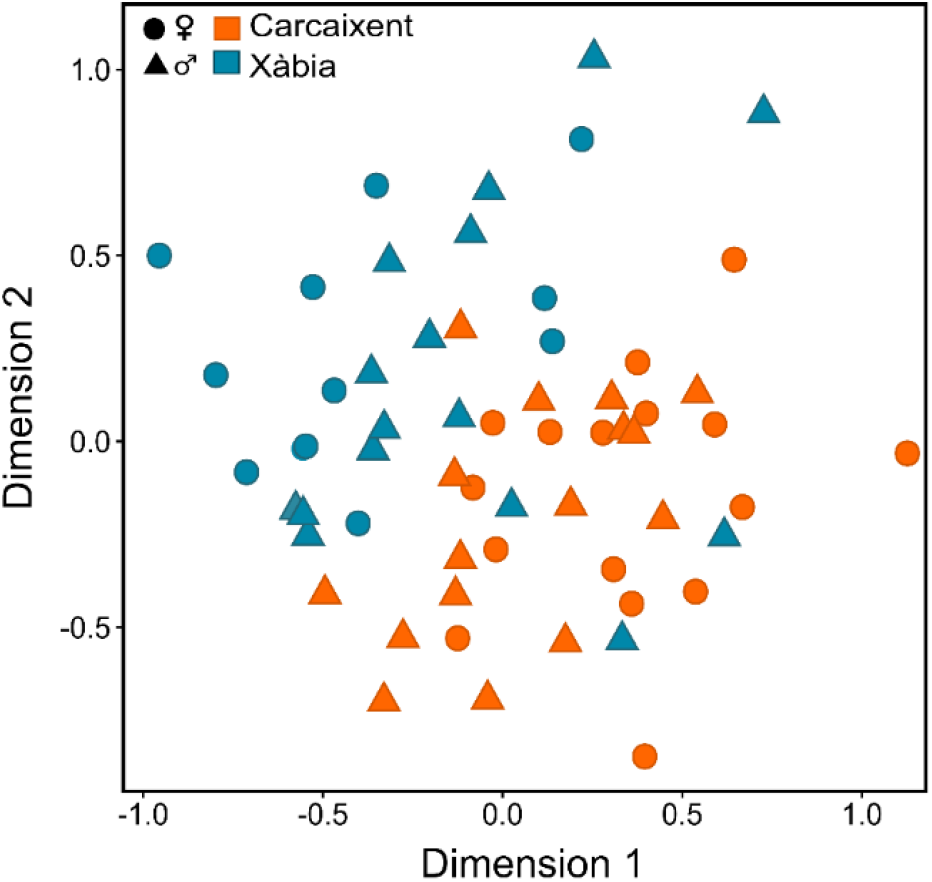
Distribution of female and male *Chamaerops humilis* plants (N= 60) in the NMDS chemospace (stress=0.24, NMDS based on VOC relative abundances) at the two study sites. Multivariate models showed that scent composition (VOC proportions) differed between the two sites (MGLM: LRT= 1889, *p*< 0.001) but not between sexes (*p*> 0.05; Appendix S1: Table S2). The results were similar for VOCs presence/absence (LRT= 235; *p*= 0.001).

For the olfactory bioassays, we first tested possible directional preferences with data from control trials by using binomial exact tests (expected value of 0.5). Then we fitted generalized linear models (GLM) with logistic regression (logit as link function, no overdispersion nor influential observations were detected) on each beetle species to test for the effect of scent origin (site) and fire treatment (burnt vs unburnt) on the binary response (VOC arm=1, control arm=0) of the beetles. We did not include plant sex in the analyses as we did not directly compare preference for scent from female and male plants in the same trials. In order to assess differences in choice proportions among VOC and control trials we performed binomial exact tests for each beetle species. To study the beetles’ preference for *C. humilis* scent vs floral scent of the co-occurring species, we used Fisher’s exact tests (for small number of trials) on VOC trials with the same co-occurring species. Significance in GLMs was tested using chi-squared Wald test with *aod* package (Lessnoff and Lancelot, 2012). All analyses were run in R software version 3.5.1 (R Core Team, 2018).

## RESULTS

### Scent composition

Leaves of *C. humilis* emitted 50 compounds (Appendix S2: Table S1), with the number of VOCs per plant ranging from 2 to 17 (7.3 ± 3.26 VOCs per sample). According to their biosynthetic origin, aliphatic (50.5% mean abundance of the total scent) were the most abundant compounds, followed by terpenoids (40.3%). The most common volatile was the monoterpene β-Ocimene (in 86% of the samples). Within aliphatic, fatty-acid hydrocarbons where the most abundant (32.5% mean abundance).

Multivariate models showed that scent composition differed between the two sites but not between sexes (Fig. 2). Fire changed overall scent composition (MGLM: burnt vs unburnt: LRT= 104.85; *p*= 0.001; for scent matrix on VOC proportions: LRT= 1066, *p*< 0.001) and its effect varied between the two localities (MGLM: interaction between site and fire treatment; LRT= 35.91; *p*= 0.04). At the two sites separately, fire also modified overall scent composition (Fig. 3). The proportions of β-Ocimene did not change after the fires (univariate tweedie GLM: Carcaixent, burnt vs unburnt: *t*= -0.25, *p*= 0.80; Xàbia, burnt vs unburnt: *t*= -0.15, *p*=0.88).

**Figure 3.**
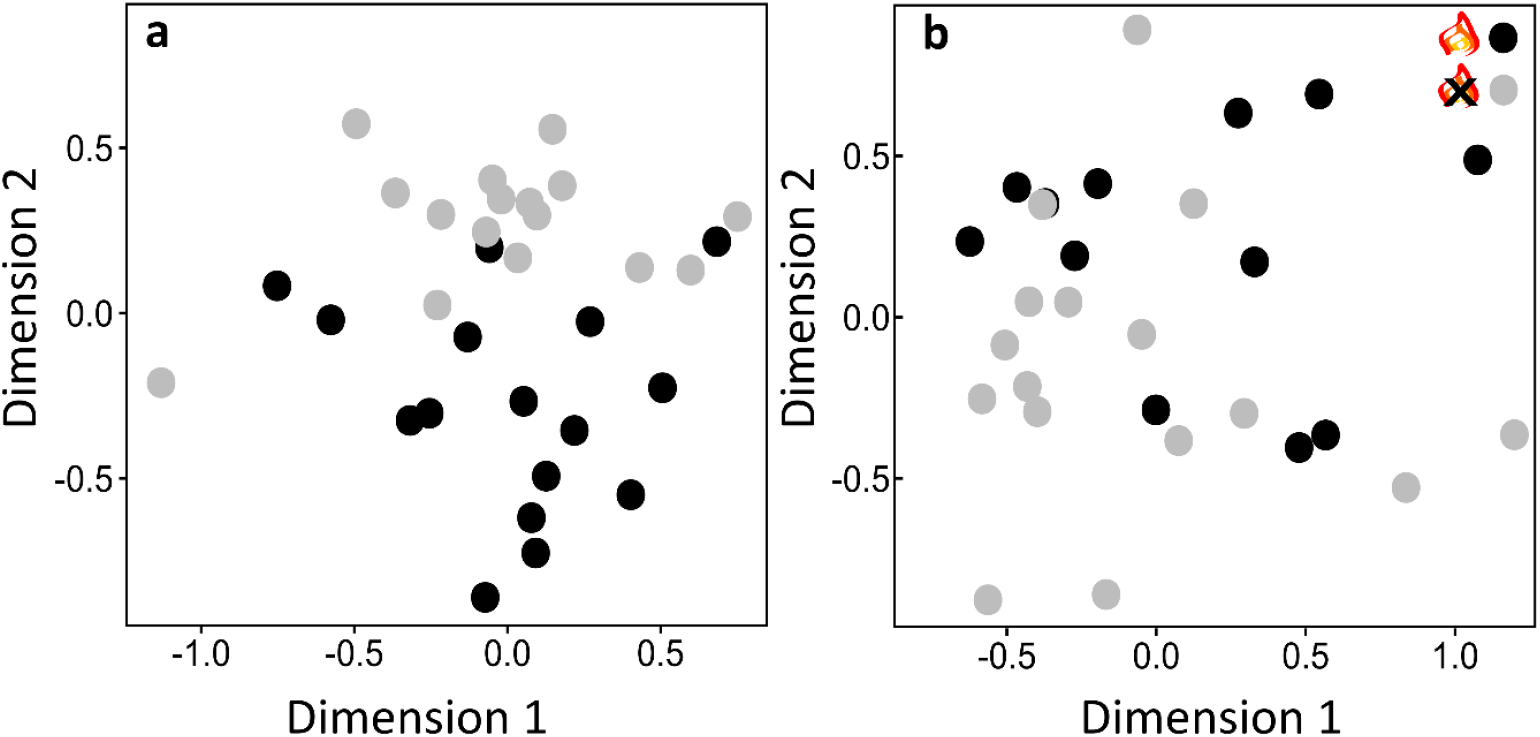
Distribution of *Chamaerops humilis* plants from burnt (black) and unburnt (grey) areas in the NMDS chemospace (based on VOC proportions; the two first dimensions are shown) for the two study sites: Carcaixent (N=28 plants, stress=0.165 two first dimensions are shown) and Xàbia (N=32 plants, stress=0.134). Scent composition differed between burnt and unburnt areas at both sites (Carcaixent, MGLM: LRT=350.1, *p=* 0.007; Xàbia, MGLM: LRT=338.4, *p*= 0.043). Results from multivariate models were similar when considering VOCs presence/absence (Carcaixent, MGLM: LRT=71.3, *p*= 0.008; Xàbia, MGLM: LRT=70.9, *p*= 0.038).

### Olfactory bioassays

The beetles showed no directional trends for any arm of the olfactometer (weevil: N= 22, *p*= 0.83; sap beetle: N= 35, *p*= 0.31). Both species displayed a clear preference for *C. humilis* scent over the control arm in VOC trials (Fig 4, Appendix S3: Videos S1, Appendix S4: Video S2). There were no differences in beetles’ choice between scents from the two study sites (logistic regression; weevil: N= 72, site χ_1_^2^= 0.43, *p*= 0.51; sap beetle: N= 65, site χ _1_^2^ =1.5, *p*= 0.22). Beetles responded in a similar way to the scent from burnt and unburnt areas (logistic regression; weevil: fire treatment χ _1_^2^ = 3.0, *p*= 0.09; sap beetle: fire treatment χ _1_^2^ = 0.15, *p*= 0.70, Fig. 4). In addition, both pollinators displayed a strong preference for *C. humilis* scent than for floral scent from most of the eight co-occurring species. Specifically, the weevil *D. chamaeropis* showed preference for *C. humilis* scent in trials with six co-occurring species and no preference in trials with *Iris sisyrinchium* and *Muscari neglectum*. The sap beetle *M. pallidulus* showed a preference for *C. humilis* in trials with five co-occurring species and no preference in trials with *Cistus salvifolius, Muscari neglectum* and *Rosmarinus officinalis* (see Appendix S1: Table S3).

**Figure 4.**
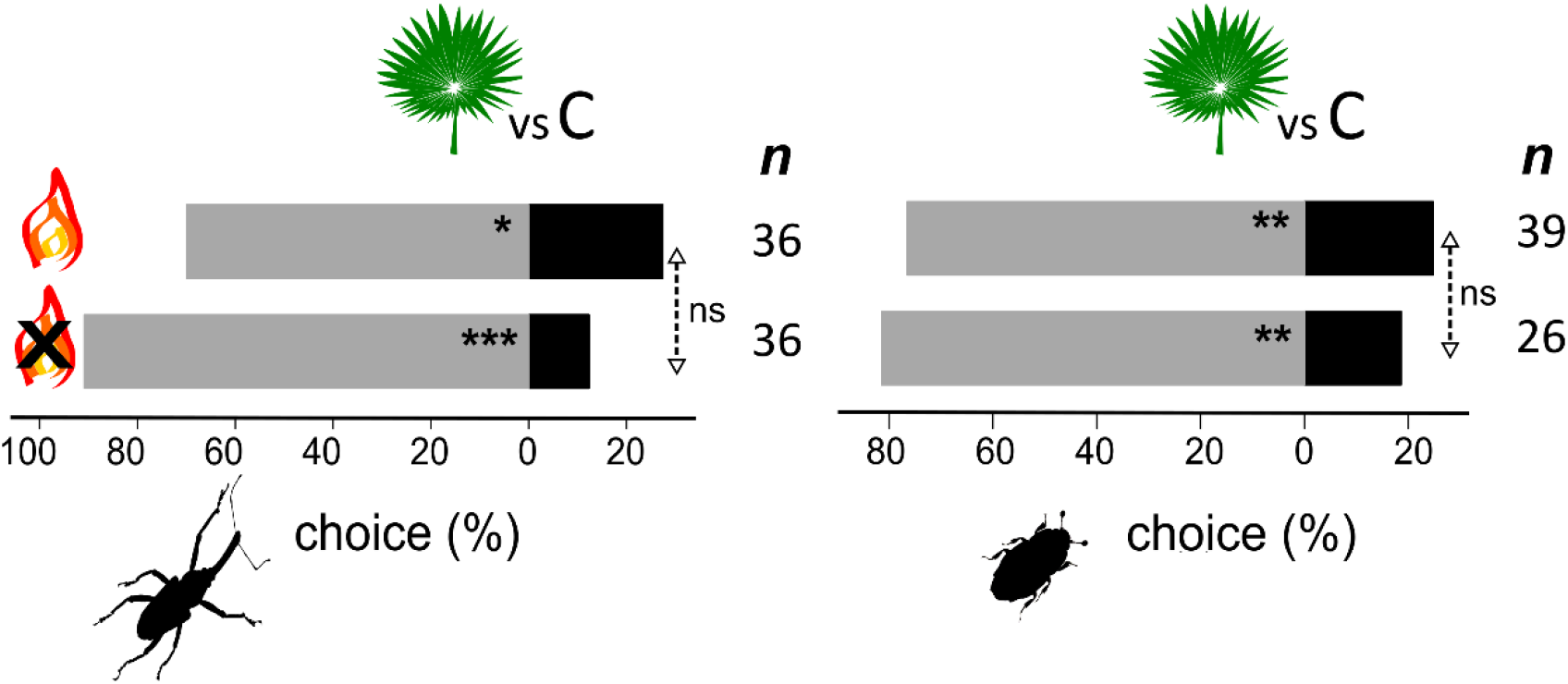
Response of the weevil *Derelomus chamaeropis* (left) and the sap beetle *Meligethinus pallidulus* (right) to *Chamaerops humilis* leaf scent in Y-tube bioassays. The grey bars refer to preference for the treatment arm with VOCs from burnt areas or unburnt areas; the black bars show response to the control arm (C: control). Significant differences between treatment and the control are denoted with asterisks (binomial tests: \**p* < 0.05; \*\**p* < 0.01; \*\*\**p* < 0.001). Dashed arrows show comparison between VOC treatments assessed by logistic GLMs (ns: not significant). *n*: number of individuals that made a choice out of 48 tested per treatment.

## DISCUSSION

Our study shows that wildfires can alter the composition of plant fragrances mediating mutualistic interactions such as pollination. In addition, and contrary to our expectation, we found a resilient response (as defined by Holling 1973) of the two pollinators that despite of changes in scent after disturbance, were similarly attracted to scent from burnt and unburnt areas. The latter adds on to previous evidence of the specialization of both pollinators in *C. humilis* and provides further insights on the role of co-pollinators in specialized systems from disturbance-prone environments.

*Chamaerops humilis* scent showed great variation among individuals and sites (see also Dufaÿ *et al*., 2004), and included common compounds in floral fragrances reinforcing the role of foliar fragrance in pollinator attraction in this system (Dufaÿ et al., 2003; 2004; Caissard et al., 2004). Fatty acid derivatives (FADs) and particularly aliphatic hydrocarbons constituted the greatest number of compounds in the scent of burnt and unburnt areas. They are frequent in fragrances involved in pollinator attraction (Knudsen et al., 2006), including other beetle-pollinated palm species (Knudsen et al., 2001) and nursery pollination systems (Bergström et al., 1991; Jürgens et al., 2002, 2003). However, the most abundant compound was the monoterpene β-Ocimene, also reported in earlier work on *C. humilis* (Dufaÿ et al., 2003; Caissard et al., 2004). Previous studies showed that β-Ocimene was more abundant in *C. humilis* scent samples collected by headspace absorption than in those from washed leaves, consistent with its function in pollinator attraction (Caissard et al., 2004). We also found the sesquiterpenoid α-Farnesene in scent samples at one our study sites, contrasting to Dufaÿ et al. (2003, 2004) that detected β-Farnesene. We detected a higher number of VOCs on *C. humilis* scent than previous work (Dufaÿ et al., 2003, 2004; Caissard et al., 2004). This could be due to our larger sample size (N=60 plants vs N=12 and 8 plants in Dufaÿ et al., 2003, 2004; Caissard et al., 2004) or, to differences in sampling (i.e. collection times, field vs greenhouse scent sampling), but also to scent variability among populations (Gaillard 2003; Caissard et al., 2004).

There is a variety of mechanisms by which fires may modify scents mediating plant communication. For instance, fire creates vegetation gaps with higher irradiance and temperature that can promote terpene emissions and modify plant blends (Farré-Armengol et al., 2014). In Mediterranean-climate regions, high intensity fires, as the ones studied here, are common, and they may alter soil moisture and nutrient availability (Keeley et al., 2011).

Drought and variation in soil nutrient content can induce changes on floral scents and pollinator attraction (Burkle and Runyon, 2016; Majetic et al., 2017; Glenny et al., 2018). In addition, antagonistic pressures modify plant fragrances (Kessler et al., 2011; Burkle and Runyon, 2016), and these pressures can be altered by wildfires (Knight and Holt, 2005; García et al., 2016; Murphy et al., 2018). At a broader scale, recent studies have pointed out the importance of plant chemical-signaling on pollination at the community level (Kantsa et al., 2017; Burkle and Runyon, 2019). In this sense, wildfires may also modify patterns of scent emissions through their effects on community structure (e.g. changes on density and diversity of plant species neighbors).

We found no evidence of changes in pollinator attraction as a response to the scent variation imposed by fire. Because not all VOCs act as attractants (Schiestl et al., 1997) and other plant cues are also involved in pollinator attraction, the observed changes in composition may not necessarily trigger a shift in pollinator behaviour (Wang et al., 2012). This lack of differential response of pollinators to the palm’s scent after fire, reinforces the idea that other factors may explain the observed postfire variation in pollinators’ abundance (García et al., 2018). This may include the variation in the life cycle of the two beetle species (tightly dependent on the inflorescences of *C. humilis* in *D. chamaeropis*, which immediately burnt after a fire), the higher densities of *M. pallidulus* in unburnt areas (potentially allowing faster re-colonization), and/or different dispersal abilities. In addition, because our results were based on differences in the frequency of beetle’s response to scent from burnt and unburnt sites, and we did not directly test for differences in their response within the same trials, we cannot completely exclude postfire changes in scent as an additional factor contributing to the reported variation in the beetle’s abundance.

From the plant’s perspective, the stability of the most abundant compounds, which remain abundant after fire, might be related to the resilience to recurrent environmental fluctuations that are common in fire-prone ecosystems. On the other hand, the persistent response of the pollinators might be explained by a potential adaptation to a dynamic odour landscape, where scent emissions frequently change in response to disturbance such as wildfires in Mediterranean landscapes (Jürgens and Bischoff, 2017). Under the hypothesis of dynamic odourscapes, organisms from ecosystems that frequently experience changes in VOC emissions may be adapted to such dynamic environments (Endler, 1992; Wilson et al., 2015; Jürgens and Bischoff, 2017). That is, fire-prone ecosystems can be viewed as dynamic odourscapes where pollinators are still responsive to fire-induced changes in scent emissions.

The experimental bioassays with co-occurring plant scents reinforce the idea of high specificity in this system, coinciding with previous field observations and pollen loads analyses (García et al., 2018). Although these results should be taken with caution because of the low number of trials with the co-flowering species, a strong specificity between plant fragrances and pollinators also occurs in other specialized systems (Proffit et al., 2009; Friberg et al., 2014). This specificity between the two pollinator species and the palm scent promotes reproductive resilience in the palm after fires, when its pollination mainly relies on the co-pollinator, and the palm shows similar fruit set levels than in the unburnt (García et al., 2018). Although co-pollinators are common in nursery-pollination systems (Thompson and Pellmyr, 1992; Prieto-Benítez et al., 2017; Hossaert-McKey et al., 2010), little research has been performed in a disturbance context where their role may be particularly relevant if nursery pollinators are sensitive to disturbance (Suchan et al., 2015; García et al., 2018).

### Concluding remarks

Researchers have only recently started to assess disturbance effects on plant scent emissions involved in pollination. Most of these studies were conducted under controlled conditions by artificially selecting the level of perturbation imposed. Here we show, for the first time, field evidence of the effects of wildfires on chemical signals mediating plant-pollinator interactions. Despite the changes detected, the interactions remained resilient to fire, even in the case of a specialized pollination system. Therefore, our results can provide an example of the Jazz hypothesis (Schmitz, 2018) in which species react to fluctuations in their environments and this flexibility may thereby impart more resilience to disturbance than previously thought.

## Acknowledgements

This research was supported by an FPI contract to YG (BES-2013-062728) and by the projects FILAS and FIROTIC (CGL2015-64086-P, PGC2018-096569-B-I00) from the Spanish Government. The authors thank Joan Nicolau-Jiménez, Guillermo Benítez-López, Beatriz López-Gurillo and Daniel Ginart-Rodriguez for their help at field. Marie-Charlotte Anstett and Samuel Prieto-Benítez provided valuable advice. We also thank Samuel Prieto-Benítez for his valuable comments on a previous version of the manuscript, Guillermo Benítez-López for his help with the set-up of the olfactory bioassays, and Lola Álvarez-Ruiz for providing the camera for video recording. The authors declare that they have no conflict of interest.

## Author contributions

YG, JGP and MCC conceived and designed the study. YG conducted the study and analysed the data. All authors interpreted the results. YG led the manuscript writing and all authors contributed to the final version.

## Data availability statement

The data supporting this article is available from the Dryad Digital Repository (https://doi.org/10.5061/dryad.2bvq83brf).

## Supplementary Information

**Appendix S1**

**Figure S1.** Y-tube olfactometer used in the experimental bioassays.

**Methods.** Details on scent collection, GC-MS analysis, olfactory bioassays procedure, and multivariate generalized linear models.

**Table S2.** Preliminary MGLM testing for differences in scent composition among *Chamaerops humilis* leaf samples.

**Table S3.** Results of Fisher’s exact tests on the olfactory bioassays with the weevil *Derelomus chamaeropis* and the sap beetle *Meligethinus pallidulus* against *Chamaerops humilis* leaf scent and floral scent of eight co-occurring species.

**Appendix S2**

**Table S1.** Volatile organic compounds detected in *Chamaerops humilis* leaf scent at burnt and unburnt areas from two localities in Spain (Carcaixent and Xàbia).

**Appendix S3**

**Video S1.** *Derelomus chamaeropis* weevil attracted by *Chamaerops humilis* scent (left arm) in the Y-tube olfactometer.

**Appendix S4**

**Video S2.** *Meligethinus pallidulus* beetle attracted by *Chamaerops humilis* scent (left arm) in the Y-tube olfactometer.

## Literature cited

Anstett, M.C. 1999. An experimental study of the interaction between the dwarf palm (Chamaerops humilis) and its floral visitor Derelomus chamaeropsis throughout the life cycle of the weevil. Acta Oecologica 20:551–558.

Bergström, G., Growth, I., Pellmyr, O., Endress, P.K., Thien, L.B., Hübener, A., and Francke, W. 1991. Chemical basis of a highly specific mutualism: chiral esters attract pollinating beetles in Eupomatiaceae. Phytochemistry 30:3221–3225.

Burkle, L.A., and Runyon, J.B. 2016. Drought and leaf herbivory influence floral volatiles and pollinator attraction. Global Change Biology 22:1644–1654.

Burkle, L.A, and Runyon, J.B. 2017. The smell of environmental change: Using floral scent to explain shifts in pollinator attraction. Applications in Plant Sciences 5:1600123.

Burkle, L.A., and Runyon, J.B. 2019. Floral volatiles structure plant–pollinator interactions in a diverse community across the growing season. Functional Ecology 33: 2116–2129.

Caissard, J.C., Meekijjironenroj, A., Baudino, S., and Anstett, M.C. 2004. Localization of production and emission of pollinator attractant on whole leaves of Chamaerops humilis (Arecaceae). American Journal of Botany 91:1190–9.

Clarke, P.J., Lawes, M.J., Midgley, J.J., Lamont, B.B., Ojeda, F., Burrows, G.E., and Knox, K.J.E. 2013. Resprouting as a key functional trait: how buds, protection and resources drive persistence after fire. New Phytologist 197:19–35.

Dixon, P. 2003. VEGAN, a package of R functions for community ecology. Journal of Vegetation Science 14:927–930.

Dötterl, S., Jahreiß, K., Jhumur, U.S., Jürgens, A. 2012. Temporal variation of flower scent in Silene otites (Caryophyllaceae): a species with a mixed pollination system. Botanical Journal of the Linnean Society 169:447–460.

Dudareva, N., and Pichersky, E. 2006. Biology of floral scent. Taylor and Francis, Boca Ratón, CRC press.

Dufaÿ, M., and Anstett, M.C. 2003. Conflicts between plants and pollinators that reproduce within inflorescences: evolutionary variations on a theme. Oikos 100:3–14.

Dufaÿ, M., and Anstett, M.C. 2004. Cheating is not always punished: killer female plants and pollination by deceit in the dwarf palm Chamaerops humilis. Journal of Evolutionary Biology 17:862–8.

Dufaÿ, M., Hossaert-McKey, M., and Anstett, M.C. 2003. When leaves act like flowers: how dwarf palms attract their pollinators. Ecology Letters 6:28–34.

Dufaÿ, M., Hossaert-McKey, M., and Anstett, M.C. 2004. Temporal and sexual variation of leaf-produced pollinator-attracting odours in the dwarf palm. Oecologia 139:392–398.

Dunn, P.K. 2017. Tweedie: Evaluation of Tweedie exponential family models. R package version 2.3.

Endler, J.A. 1992. Signals, signal conditions, and the direction of evolution. The American Naturalist 139: S125–S153.

Farré-Armengol, G., Filella, I., Llusià, J., and Peñuelas, J. 2013. Floral volatile organic compounds: between attraction and deterrence of visitors under global change. Perspectives in Plant Ecology, Evolution and Systematics 15:56–67.

Farré-Armengol, G., Filella, I., Llusià, J., Niinemets, Ü., and Peñuelas, J. 2014. Changes in floral bouquets from compound-specific responses to increasing temperatures. Global Change Biology 20:3660–3669.

Farré-Armengol, G., Peñuelas, J., Li, T., Yli-Pirilä, P., Filella, I., Llusià, J., and Blande, J.D. 2016. Ozone degrades floral scent and reduces pollinator attraction to flowers. New Phytologist 209:152–160.

Friberg, M., Schwind, C., Roark, L.C., Raguso, R.A., and Thompson, J.N. 2014. Floral scent contributes to interaction specificity in coevolving plants and their insect pollinators. Journal of Chemical Ecology 40:955–965.

Gaillard, A. 2003. Variabilité geographique de l’attraction dans le mutualisme obligatoire Palmier nain/ charançon. DEA Biologie de l’Evolution et Ecologie, Université Montpellier II, Montpellier, France.

García, Y., Castellanos, M.C. and Pausas, J.G., 2016. Fires can benefit plants by disrupting antagonistic interactions. Oecologia, 182:1165–1173.

García, Y., Castellanos, M.C., and Pausas, J.G. 2018. Differential pollinator response un-derlies plant reproductive resilience after fires. Annals of Botany 122:961–971

Giner, G., and Smyth, G.K. 2016. Statmod: probability calculations for the inverse Gaussian distribution. R Journal 8:339–351.

Glenny, W.R., Runyon, J.B., and Burkle, L.A. 2018. Drought and increased CO2 alter floral visual and olfactory traits with context-dependent effects on pollinator visitation. New Phytologist 220:785–798.

Holling, C.S. 1973. Resilience and stability of ecological systems. Annual Review of Ecology and Systematics, 1:1–23.

Hossaert-McKey, M., Soler, C., Schatz, B., and Proffit, M. 2010. Floral scents: their roles in nursery pollination mutualisms. Chemoecology 20:75–88.

Jürgens, A., Witt, T., and Gottsberger, G. 2002. Flower scent composition in night-flowering Silene species (Caryophyllaceae). Biochemical Systematics and Ecology 30:383–397.

Jürgens, A., Witt, T., and Gottsberger, G. 2003. Flower scent composition in Dianthus and Saponaria species (Caryophyllaceae) and its relevance for pollination biology and taxonomy. Biochemical Systematics and Ecology 31:345–357.

Jürgens, A., Bischoff, M. 2017. Changing odour landscapes: The effect of anthropogenic volatile pollutants on plant-pollinator olfactory communication. Functional Ecology 31:56–64.

Kantsa, A., Raguso, R.A., Dyer, A.G., Sgardelis, S.P., Olesen, J.M., and Petanidou, T. 2017. Commnity-wide integration of floral colour and scent in a Mediterranean scrubland. Nature Ecology and Evolution 1:1502.

Keeley, J.E., Bond, W.J., Bradstock, R.A., Pausas, J.G. and Rundel, P.W., 2011. Fire in Mediterranean ecosystems: ecology, evolution and management. Cambridge University Press.

Kessler, A., Halitschke, R., Poveda, K. 2011. Herbivory-mediated pollinator limitation: negative impacts of induced volatiles on plant-pollinator interactions. Ecology 92:1769–1780.

Knight, T.M. and Holt, R.D., 2005. Fire generates spatial gradients in herbivory: an example from a Florida sandhill ecosystem. Ecology, 86:587–593.

Knudsen, J., Tollsten, L., and Ervik, F. 2001. Flower scent and pollination in selected neotropical palms. Plant Biology 3:642–653.

Knudsen, J.T, Eriksson, R, Gershenzon, J, and Ståhl, B. 2006. Diversity and distribution of floral scent. The Botanical Review 72:1–120.

Lesnoff, M., and Lancelot, R. 2012. aod: Analysis of Overdispersed Data. R package version 1.3.1, URL http://cran.r-project.org/package=aod

Li. T, Blande, J.D., and Holopainen, J.K. 2016. Atmospheric transformation of plant volatiles disrupts host plant finding. Scientific Reports 6:33851.

Majetic, C.J., Fetters, A.M., Beck, O.M., Stachnik, E.F., and Beam, K.M. 2017. Petunia floral trait plasticity in response to soil nitrogen content and subsequent impacts on insect visitation. Flora 232:183–193.

Murphy, S.M., Vidal, M.C., Smith, T.P., Hallagan, C.J., Broder, E.D., Rowland, D. and Cepero, L.C., 2018. Forest fire severity affects host plant quality and insect herbivore damage. Frontiers in Ecology and Evolution, 35.

Nicholson, C.C., and Egan, P.A. 2019. Natural hazard threats to pollinators and pollination. Global Change Biology. doi: 10.1111/gcb.14840

Pausas, J.G., and Fernández-Muñoz, S. 2012. Fire regime changes in the Western Mediterranean Basin: from fuel-limited to drought-driven fire regime. Climatic Change 110:215–226.

Prieto-Benítez, S., Dötterl, S., & Giménez-Benavides, L. 2015. Diel variation in flower scent reveals poor consistency of diurnal and nocturnal pollination syndromes in Sileneae. Journal of Chemical Ecology 41: 1095–1104.

Prieto-Benítez, S., Dötterl, S., & Giménez-Benavides, L. 2016. Circadian rhythm of a Silene species favours nocturnal pollination and constrains diurnal visitation. Annals of Botany 118: 907–918.

Prieto-Benítez, S., Yela, J.L., and Giménez-Benavides, L. 2017. Ten years of progress in the study of Hadena-Caryophyllaceae nursery pollination. A review in light of new Mediterranean data. Flora 232:63–72.

Proffit, M., Chen, C., Soler, C., Bessière, J.M., Schatz, B., and Hossaert-McKey, M. 2009. Can chemical signals, responsible for mutualistic partner encounter, promote the specific exploitation of nursery pollination mutualisms? The case of figs and fig wasps. Entomologia Experimentalis et Applicata 131:46–57.

R Core Team. 2018. R: A language and environment for statistical computing. R Foundation for Statistical Computing, Vienna, Austria. http://www.R-project.org/

Schiestl, F.P., Ayasse, M., Paulus, H.F., Erdmann, D., and Francke, W. 1997. Variation of floral scent emission and postpollination changes in individual flowers of Ophrys sphegodes subsp. sphegodes. Journal of Chemical Ecology 23:2881–2895.

Schlumpberger, B.O, and Raguso, R.A. 2008. Geographic variation in floral scent of Echinopsis ancistrophora (Cactaceae); evidence for constraints on hawkmoth attraction. Oikos 117: 801–814.

Schmitz, O.J. 2018. Species in ecosystems and all that jazz. PLOS Biology, 16: e2006285.

Suchan, T., Beauverd, M., Trim, N., and Alvarez, N. 2015. Asymmetrical nature of the Trollius–Chiastocheta interaction: insights into the evolution of nursery pollination systems. Ecology and Evolution 5:4766–4777.

Swab, R.M., Regan, H.M., Keith, D., Regan, T.J., Ooi, M.K. 2012. Niche models tell half the story: spatial context and life-history traits influence species responses to global change. Journal of Biogeography 39:1266–1277.

Tavsanoglu, Ç, and Pausas, J.G. 2018. A functional trait database for Mediterranean Basin plants. Scientific Data 5:180135.

Thompson, J.N., and Pellmyr, O. 1992. Mutualism with pollinating seed parasites amid copollinators: constraints on specialization. Ecology 73:1780–1791.

Wang, G., Compton, S.G., and Chen, J. 2012. The mechanism of pollinator specificity between two sympatric fig varieties: a combination of olfactory signals and contact cues. Annals of Botany 111:173–181.

Wang, Y., Naumann, U., Wright, S.T., Eddelbuettel, D., and Warton, D.I. 2017. mvabund: Statistical methods for analysing multivariate abundance data. R package version 3.12.3.

Westerling, A.L., Hidalgo, H.G., Cayan, D.R., Swetnam, T.W. 2006. Warming and earlier spring increase western US forest wildfire activity. Science 313:940–943.

Wilson, J.K., Kessler, A., and Woods, H.A. 2015. Noisy communication via airborne infochemicals. BioScience 65:667–677.

