## Appendix_S1 for "Wildfires change plant scents but not pollinator attraction in a Mediterranean palm"

**Figure S1.** Y-tube olfactometer used in the experimental bioassays (Vidrafoc ®, Valencia Spain). The tube inner diameter was 1.5 cm, the basal branch was 8 cm long and each of the Y-branches 6 cm long. Compressed air entered at a constant flow rate through a glass flask with activated charcoal connected with a second flask with distilled water. Teflon tubes were used to connect the system (Labbox, Barcelona Spain).

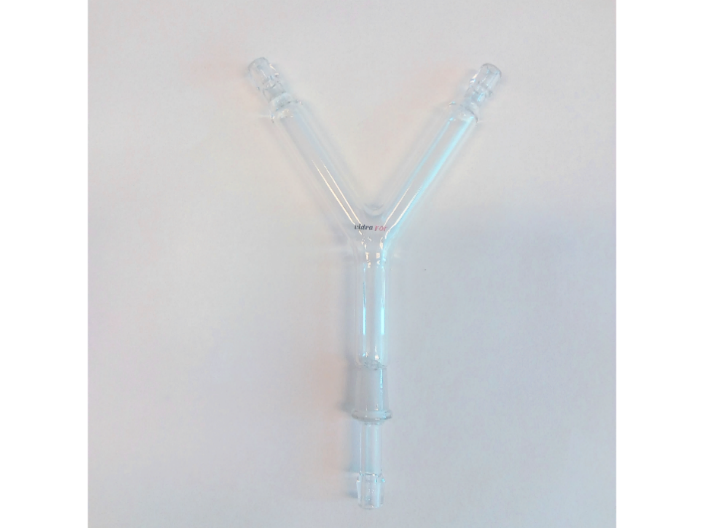

**Methods**

Details on scent collection, GC-MS analysis, olfactory bioassays and multivariate generalized models.

*Scent collection*

The scent traps were made with modified Teflon tubes (id= 6 mm, Labbox, Barcelona, Spain) previously washed with acetone and dried. Each trap contained a mixture 1:1 of 5 mg Tenax-TA (mesh 20-40) and 5 mg Carbotrap (mesh 60-80) adsorbents (Supelco® Sigma Aldrich, St. Louis, USA) between silanized glass wool (Panreac, Applichem). Samples were stored at -20 ºC before analysis and then eluted with 200 µl of dichloromethane (Sigma Aldrich) for the GC-MS analysis. For the olfactory bioassays, the leaf scent from four additional *C. humilis* individuals per sex at burnt and unburnt areas was sampled during 10 minutes in scent traps filled with a mixture 1:1 of 25 mg of each adsorbent.

To assess the specificity between the palm chemical signal and the beetle pollinators we collected the scent of flowers from eight co-occurring plant species. Seven of these species were flowering at the same time of *C. humilis* in the study sites, while *Iris* *sisyrinchium* flowered earlier. In this case, one inflorescence or five flowers per individual (for species with solitary flowers) of each plant species were collected in the field and stored at -20ºC. The scent from two individuals of each species was collected by diluting the flowers in 1ml high-grade acetone (Chromasolv ®) and stored at 4 ºC previous to the olfactory bioassays. Leaf VOCs samples from *C. humilis* inside the scent traps were diluted with high-grade acetone and stored in the same conditions.

*GCMS analysis*

The analysis of the leaf scent was carried out by the Mass Spectrometry Section of The Experimental Research Support Service (SCSIE) of the University of Valencia using an Agilent 7890B gas chromatograph (GC) coupled with a Agilent 5977A mass spectrometer (MS) and separated on a HP-5 MS capillary column (30 m x 0.25 mm inside diameter, 0.25 µm film thickness). Helium was used as the carried gas. Oven temperature was held at 60 ºC for 5 min and increased by 5 ºC min^-1^ until 180 ºC, and then by 25 ºC min^-1^ until 280 ºC under a flow of 1.1 ml min^-1^.

In order to build the scent data matrix, contaminant VOCs present in the ambient controls were corrected from scent samples. Several compounds in scent samples not known as floral nor foliar volatiles (*Pherobase* semiochemical data base, El-Sayed 2012) and with high molecular weight were also removed (e.g. a phenyl ester of pentanoic acid and ethylhexyl benzenedicarboxylic acid ester).

*Olfactory bioassays*

The olfactory bioassays started 48 hours after pollinator sampling in the field and lasted 9 days. We performed a series of pilot tests in the lab with the two beetles (the two species separately and one individual per assay) and three different scent volumes (10 µl, 20 µl and 50 µl). Both species responded (in 3’ minute tests) to the lowest amount so we chose 10 µl as test amount in the bioassays. At each trial, the response of one beetle individual was monitored for 3 min, and the choice was recorded when a beetle walked to the end of one of the arms and remained there for more than 10 seconds. We ran 384 trials ([24 with VOCs + 24 controls] x 2 plant sex x 2 beetle species x 2 study sites). Because of the high number of trials and the difficulty to maintain the beetles at the laboratory, 187 beetles were tested twice (95 *D. chamaeropis* and 92 *M. pallidulus*). The second trials were always performed on a different day of the first trials for this beetles’ subset. We did not detected differences in response between beetles tested one or two times (logistic GLMs on each beetle species responses and factor “test” with levels: once or twice as predictor; *p*> 0.05 in both cases). The olfactometer and the glass flasks were washed with an odourless detergent and acetone and dried between trials. All bioassays were conducted under red light conditions (20W-1000 Lumen-1P66, Matel ®) to avoid visual cues.

*Olfactory bioassays with co-occurring plant species*

For the olfactory bioassays with *C. humilis* scent and floral scent of the co-occurring plants, we used on each trial a 10µl sample of floral VOCs in one arm of the olfactometer and in the other arm a 10µl sample from *C. humilis* leaf VOCs, both diluted in acetone. Eight VOC trials plus eight alternated control trials (i.e. only acetone) were conducted with each beetle pollinator for each of the co-occurring plant species. We followed the same procedure to assess the response of each beetle individual as on trials with scent from the burnt and unburnt areas.

*Multivariate generalized linear models (MGLMs)*

The MGLM approach fits a GLM to each response variable with a common set of predictors and uses a resampling method of rows of the data matrix. To test for differences in the estimated multivariate response (i.e. overall scent composition), the method uses ANOVA (likelihood ratio test) with 999 bootstrap interactions by probability integral transform (PIT-trap) residuals (Wang et al. 2012). Compared to distance-based approaches

(PERMANOVAs, SIMPER), MGLMs are robust at detecting differences in the effects of

location and dispersion in multivariate data sets and allow to include a mean-variance relationship (Wang et al. 2012; Warton et al. 2012).

**References**

El-Sayed AM (2012) The Pherobase: database of pheromones and semiochemicals. In: The

Pherobase. http://www.pherobase.com.

Wang Y, Naumann U, Wright ST, Warton DI (2012) Mvabund-an R package for model-based analysis of multivariate abundance data. Methods Ecol Evol 3:471–474.

Warton DI, Wright ST, Wang Y (2012) Distance-based multivariate analyses confound location and dispersion effects. Methods Ecol Evol 3:89-101.

**Table S2.** Preliminary MGLM testing for differences in scent composition among *Chamaerops humilis* leaf samples (N=60) according to study site (Carcaixent vs. Xàbia), plant sex (female vs. male plants), fire treatment (burnt vs. unburnt areas), stem length and the interaction between site and fire treatment. Df _res, diff_ = Degrees of freedom. Significant differences are in bold.

| **Response** | **Predictors** | **Df _res, diff_** | **Deviance** | ***P*-value** |
| --- | --- | --- | --- | --- |
| Scent composition | Site | 58, 1 | 235.03 | **0.001** |
|  | Fire treatment | 57, 1 | 104.85 | **0.002** |
|  | Plant sex | 56, 1 | 65.03 | 0.185 |
|  | Stem length | 55,1 | 36.21 | 0.969 |
|  | Site*Fire treatment | 54, 1 | 43.61 | **0.033** |

**Table S3.** Results of Fisher’s exact tests on the olfactory bioassays with the weevil *Derelomus chamaeropis* and the sap beetle *Meligethinus pallidulus* against *Chamaerops humilis* leaf scent and floral scent of eight co-occurring species from the study sites. Significant *P*-values are in bold and correspond with a higher preference for *C. humilis* scent in the Y-tube olfactometer. N= number of trials out of 8 where the beetles’ preferred *C. humilis* scent.

|  |  |  | **Pollinator species** | | | |
| --- | --- | --- | --- | --- | --- | --- |
| **Plant species** |  | *D. chamaeropis* | |  | *M. pallidulus* | |
|  | N | d.f | *P*-value | N | d.f | *P*-value |
| *C. humilis* vs *Cistus salvifolius* L. | 6 | 1 | **0.04** | 3 | 1 | n.s |
| *C. humilis* vs *Genista scorpius* (L.) DC. | 5 | 1 | **0.02** | 6 | 1 | **0.01** |
| *C. humilis* vs *Gladiolus illyricus* W.D.J. Koch | 7 | 1 | **0.01** | 5 | 1 | **0.04** |
| *C. humilis* vs *Iris sisyrinchium* (L.) Parl | 5 | 1 | n.s | 5 | 1 | **0.04** |
| *C. humilis* vs *Lavandula angustifolia* Mill. | 5 | 1 | **0.04** | 5 | 1 | **0.02** |
| *C. humilis* vs *Minuartia hybrida* (Vill.) Schischk | 8 | 1 | **0.001** | 7 | 1 | **0.01** |
| *C. humilis* vs *Muscari neglectum* Guss. Ex Ten. | 4 | 1 | n.s | 2 | 1 | n.s |
| *C. humilis* vs *Rosmarinus officinalis* L. | 5 | 1 | **0.02** | 4 | 1 | n.s |
